# Distinct Hippocampal Neuronal Reactions Reveal Different Neuronal Codes for Memory Generalization

**DOI:** 10.1101/2021.06.24.449806

**Authors:** Jun Guo, Duc Truong, Andrea Barreiro, Da-Ting Lin, Wei Xu

## Abstract

To survive in an ever-changing world we need to learn and memorize associations of environmental stimuli and generalize them to new situations. Both memory and generalization critically rely on the hippocampus, but it is unclear how hippocampal neuronal activities represent memory and generalization, and if a conserved hippocampal mechanism serves these functions. Here we compared neuronal activities in hippocampal CA1 region of two sub-strains of the widely used C57BL/6 mice, C57BL/6J (B6J) and C57BL/6NCrl (B6NCrl), in contextual fear conditioning. Both sub-strains learnt well but differed in freezing and generalization. They displayed distinct early-late bi-phasic reactions to the unconditioned stimulus. While in both sub-strains the neurons showing late-phase reactions were preferentially engaged in memory representation, the neuronal activity feature that correlated with generalization level differed in the two sub-strains: in B6NCrl, these neurons’ activity level during learning negatively correlated with the generalization level; in B6J, functional coupling of these late-phase neurons with other neurons positively correlated with the generalization level. We further found that the distinct neuronal reactions were accompanied by distinct GABAb receptor-mediated inhibition but not by differences in the major synaptic inputs or neuronal excitability of the CA1. Therefore, this comparative study reveals two signature neuronal activity features in learning that can predict generalization levels. The results also demonstrate that differences in hippocampal network properties lead to diverse hippocampal mechanisms in memory encoding and generalization.

## Introduction

We learn from experience to handle new situations. A basic type of learning and memory is conditioning, in which conditioned stimulus (CS) becomes associated with unconditioned stimulus (US) and this association expands our behavioral repertoire (Fanselow and Poulos, 2005; Maren, 2001). Since the world continuously changes and we may never encounter an identical situation again, the associations established through conditioning need to be generalized to similar situations to be useful (Shepard, 1987). Together, conditioning and generalization build the foundation for our adaptive behavior. The hippocampus is well-known for its central roles in memory (Buzsaki and Moser, 2013; Eichenbaum, 2004; Eichenbaum et al., 1987; Fried et al., 1997; Sakurai, 2002). The hippocampus is critically involved in generalization as well (Asok et al., 2018; Berens and Bird, 2017; Kumaran and McClelland, 2012). Functional manipulations of the hippocampus in model animals or pathological damage to human hippocampus lead to various deficits in memory and generalization (Jeneson and Squire, 2012; Tonegawa et al., 2015; Xu and Sudhof, 2013). Recently multiple-unit recording and *in vivo* imaging techniques have revealed the neuronal activities in the hippocampus in studies related to learning and memory, including contextual fear conditioning, a popular animal model for studying learning, memory and generalization (Jimenez et al., 2020; Moita et al., 2004; Munera et al., 2001; Tanaka et al., 2018; Wu et al., 2017). Despite significant progress, we are still far from having a clear mechanism for learning, especially generalization.

The hippocampus has been studied in different model systems, including mice, rats, non-human primates and human subjects. The anatomical organization of the hippocampal circuits and the physiological properties of the neurons making up these circuits are highly stereotypical across these mammals (van Strien et al., 2009). This similarity suggests that hippocampus may operate with a conserved neuronal and circuitry mechanism. Although this “conserved” view of hippocampal functions is supported by some studies (Bunsey and Eichenbaum, 1996), there are considerable differences across species in hippocampal structure, molecular contents, physiological properties and hippocampus-dependent behaviors (Bingman, 1992; Koopmans et al., 2018; McNamara et al., 1996; West, 1990). It therefore remains unclear how conserved and diverse the hippocampal neuronal representation of memory and generalization can be.

To identify the neuronal activities linked to conditioning and generalization, and to test how conserved / diverse hippocampal information processing is, we conducted functional imaging of hippocampal neuronal activity in two sub-strains of C57BL/6 mice, which are almost identical in genetic backgrounds but differ significantly in memory and generalization. C57BL/6J (B6J) mouse line has been established and maintained in JAX since 1937; but diverged to multiple sub-strains including C57BL/6NCrl (B6NCrl). These two sub-strains—B6J and B6NCrl – differ greatly in the contextual fear conditioning. Both sub-strains learn well but exhibit distinct levels of fear responses and generalization (Radulovic et al., 1998; Stiedl et al., 1999). This allows us to examine how conserved the hippocampal information processing is in animals with highly similar genetic backgrounds. We found that the hippocampal CA1 regions in the sub-strains showed highly different neuronal reactions to the US as well as distinct activity patterns that could predict fear generalization levels. Our data indicate diverse hippocampal neuronal processing in the two closely related mouse sub-strains. We further demonstrated that this difference may arise from the different feedforward inhibitions in the hippocampus.

## Result

### Imaging hippocampal neuronal activity in mice with distinct memory and generalization

We first characterized the behavior of B6J and B6NCrl mice in contextual fear conditioning to examine the acquisition, retrieval and generalization of memories. The mice were trained in a novel behavioral cage (Context). They received footshocks as an unconditioned stimulus (US). To examine fear memory and its generalization, on the second day, the mice were tested either in the original training context, or in a context with one or multiple sensory cues altered (Fig. 1A). Time spent on freezing was measured as an indicator of fear memory. Compared to B6J, B6NCrl mice showed faster learning during training (Fig. 1B), and higher freezing levels in both the original context and the contexts with altered sensory cues (Fig. 1A, 1C). Consistent with earlier studies, B6J mice showed a steeper generalization gradient (Fig. 1C).

**Fig. 1:**
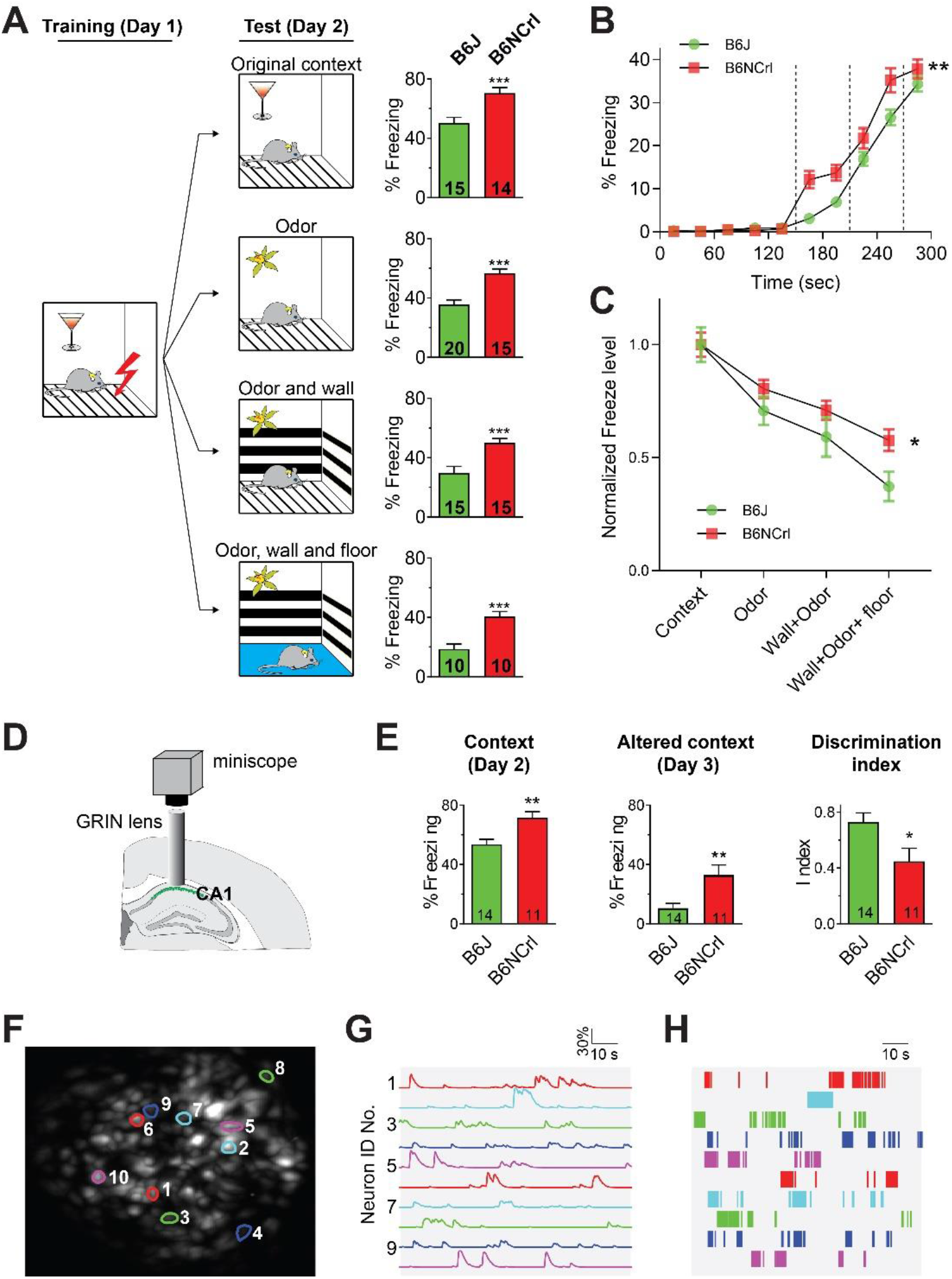
Functional Imaging of mice with distinct learning and memory. **(A)** Behavioral measurement of contextual fear memory and memory generalization in B6J and B6NCrl. The mice from both sub-strains were trained for contextual fear conditioning in a novel context on Day 1. After training, mice were randomly separated into 4 groups and each group was tested in one of the 4 contexts with increasing differences from the original training context on Day 2. B6NCrl mice froze significantly more than B6J mice in all the contexts tested. The numbers in the bars indicate the numbers of mice been tested. (*** P<0.001, Student’s t-test) **(B)** Faster learning in B6NCrl vs. B6J mice. Freezing level during fear conditioning training were plotted in 30-second bins. The dashed lines indicate the time when the footshocks were delivered. (n= 80 B6J mice and 39 B6NCrl mice, ** P<0.01, Two-Way ANOVA) **(C)** Generalization of contextual fear memories in B6J vs. B6NCrl mice. The freezing level was normalized to that in original context. B6J mice had steeper generalization gradient compared to that of B6NCrl (* P<0.05, Two-Way ANOVA). **(D)** Schematics showing functional imaging of CA1 with miniscope through implanted GRIN lens. Calcium indicator GCaMP6s was expressed in CA1 by local injection of AAV. The GRIN lens was implanted into the pyramidal layer of the CA1. **(E)** Behavioral performance of the mice that underwent imaging in contextual fear conditioning and tests. The numbers in the bars indicate the numbers of mice been tested. (**P<0.01, * P<0.05, Student’s t-test) **(F)** Maximum projection of the images collected in a miniscope imaging session. **(G)** Calcium signals (dF/F) of the neurons with corresponding ID No. in **F**. **(H)** Calcium signals (dF/F) shown in **G** were deconvoluted into calcium events.

To understand the neural mechanisms for the behavioral differences, we performed *in vivo* calcium imaging of hippocampal neuronal activity with miniscopes (Fig. 1D). We expressed calcium indicator GCaMP6s in the hippocampus by injecting AAV, and implanted a GRIN lens for miniscope imaging into the CA1 region of the dorsal hippocampus, a region critically involved in memory encoding and retrieval. Mice were trained and tested with a three-day contextual fear conditioning protocol. After training on Day 1, mice were tested in the original context (Context) on Day 2 and an altered context on Day 3. In the altered context, the background odor, the decoration of the walls and the floor were different from the original context. Discrimination index measured the generalization level: a higher discrimination index indicates less generalized memory [discrimination index = (freezing level in Context - freezing level in Altered context test) / (freezing level in Context + freezing level in Altered context)]. Compared to B6J, B6NCrl mice again showed higher freezing to Context and Altered context, and demonstrated a higher level of generalization, indicating that the imaging procedures did not alter the behavior of the mice (Fig. 1E). 14 B6J and 11 B6NCrl mice were imaged during fear conditioning training or testing. The calcium signal from individual neurons was extracted by CNMF-E method (Zhou et al., 2018) and deconvoluted to calcium event for further analysis (Fig. 1F-H). 57-249 neurons were recorded from each animal.

### Distinct hippocampal reactions to the US

We then compared neuronal activities of the two sub-strains in training (Fig. 2). The neuronal activities of two representative mice for B6J and B6NCrl, respectively, are shown in the raster plots in Fig.2A. To reveal the temporal profile of these activities, we averaged the neuronal activity within 2-second bins, yielding binned event rates of 150 bins for a 300-s training session. Then the activity was averaged across all mice within a sub-strain to generate an averaged event rate (Fig. 2A, middle panels). The averaged or peak neuronal activities in the different stages of the training, including the first two minutes of exploration period, the tones and the post-footshock periods were also calculated (Fig. 2B-D)

**Fig. 2:**
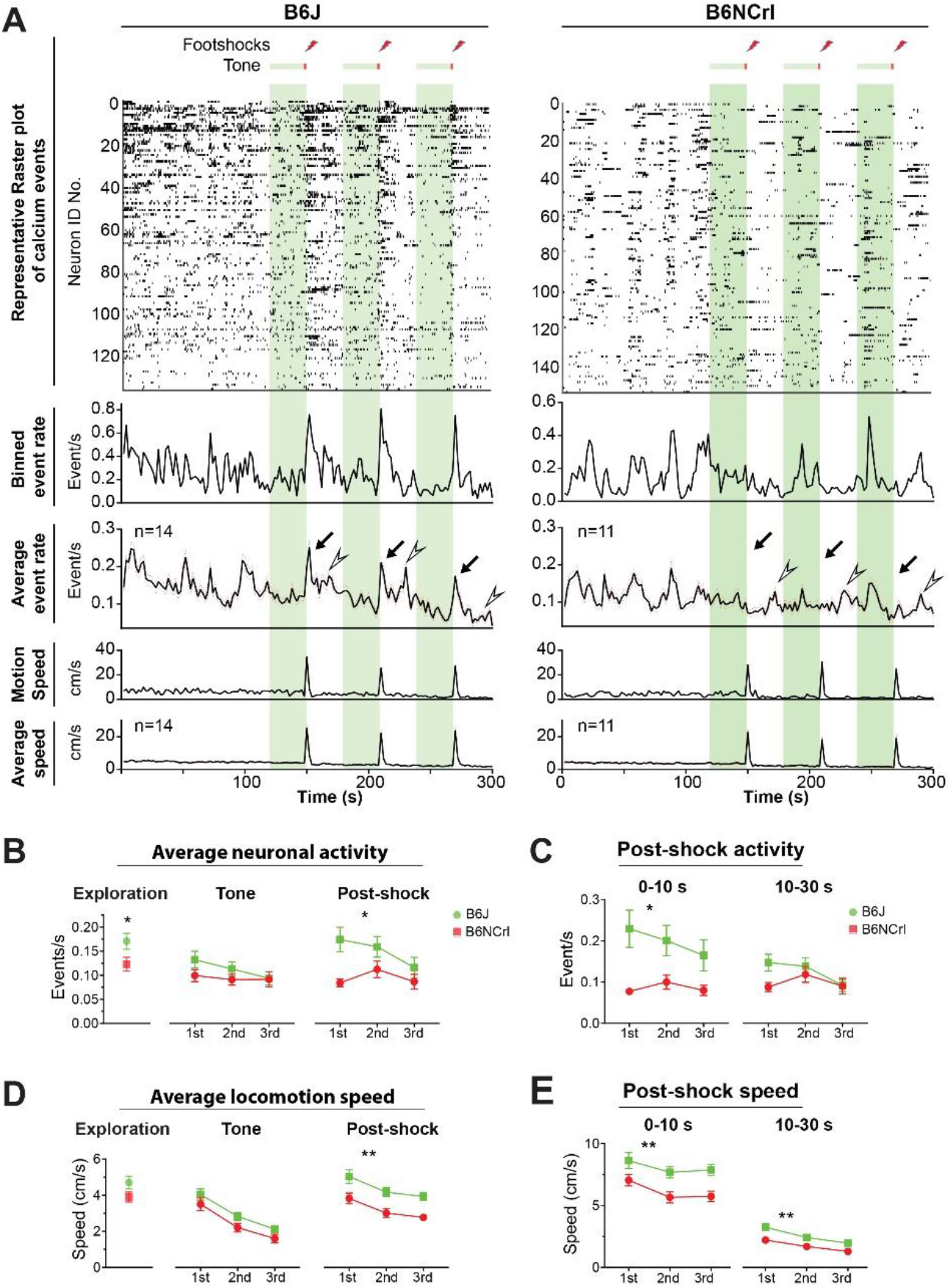
Distinct hippocampal reactions to the US. **(A)** The CA1 neuronal activities and animal locomotor activities of B6J and B6NCrl mice during contextual fear conditioning training. From top to bottom: raster plot of the neuronal activity of representative mice; the binned event rate measured with 2-s bins for the representative mice; the average event rate for the mice recorded in each sub-strain (N=14 B6J and 11 B6NCrl mice); the speed of the representative mice, and the averaged speed for the animals in each sub-strain. For the average event rate, the data are mean ± SEM (standard error of the mean, shown by orange dots). The black arrows and white arrowheads indicate the immediate or delayed reactions of the hippocampus to the US, respectively. **(B, C)** Quantification of neuronal activity in different time periods during training. (* P<0.05, Student t-test for “Exploration”; Two-way ANOVA for “Post-shock”.) **(D, E)** Quantification of mice’s locomotion in different time periods during training. (P<0.01, Two-way ANOVA.)

CA1 neuronal activities in both sub-strains fluctuated during training. Neurons in B6J showed higher overall activity both before and after the mice received the US (Fig. 2A and 2B-D). A striking difference between the two sub-strains was their reaction to the US: in B6J mice CA1 neuronal activity increased dramatically upon and right after the delivery of each of the US, whereas these increases were minimal in B6NCrl mice (Fig.2A, 2C and Supplementary video-1). In B6J this early-phase reaction was followed by a second wave of activation 10-30 seconds after the US. The late-phase activation had a lower amplitude in averaged event rates compared to the early-phase reaction. Similar to B6J, B6NCrl mice also showed the late-phase reaction(Fig 2A).

CA1 neuronal activities often correlate with animals’ locomotion (Geiller et al., 2020). To test if the different neuronal activities were related to animal behavior, we measured the locomotion speed of mice and averaged it within a 2-s bin to align it with neuronal activity (Fig. 2A, lower panels). B6J mice showed higher locomotion in the exploration stage, consistent with the higher average neuronal activities in this period. Upon the delivery of the US, both B6J and B6NCrl mice displayed a massive increase in locomotion speed, indicating that the reduction of the early-phase response to the US in B6NCrl was not due to a lack of behavioral responses (Fig 2A).

### Neurons with delayed reactions play special roles in memory encoding and generalization

We then analyzed the neuronal activities in training together with those in recall (context test in Day 2) or generalization (altered context test in Day 3) (Fig.3). The sharp contrast between B6J and B6NCrl in neuronal reactions to the US suggests that the two sub-strains may represent contextual information (the conditioned stimulus, CS) and the US differently. To reveal the potential function of US-activated neurons, we put neurons into 4 groups based on the time window in which they show increased activities (Fig. 3A). Each neuron’s activity during 0-10 s or 10-30 s from the onset of shock was compared with that in the −10 to 0 s time window with a shuffling approach (see details in Method). The neurons showing significantly higher activity in 0-10 s time window were defined as immediate response neurons (IR). The neurons with significantly higher activity during 10-30 s time window were defined as delayed response neurons (DR). Neurons showing significantly higher activity in both time windows were defined as continuous responding neurons (Co). The rest of neurons were non-responding neurons (NR). The averaged neuronal activities of the 4 groups showed higher amplitudes in the corresponding time windows, indicating the effectiveness of the shuffling approach in classifying these neurons (Fig. 3B). Consistent with the pronounced reaction to the US, B6J mice had a higher percentage of neurons in IR group than B6NCrl (510/2139 vs 197/1668, p<0.001, Chi-square test) (Fig. 3C).

**Fig. 3:**
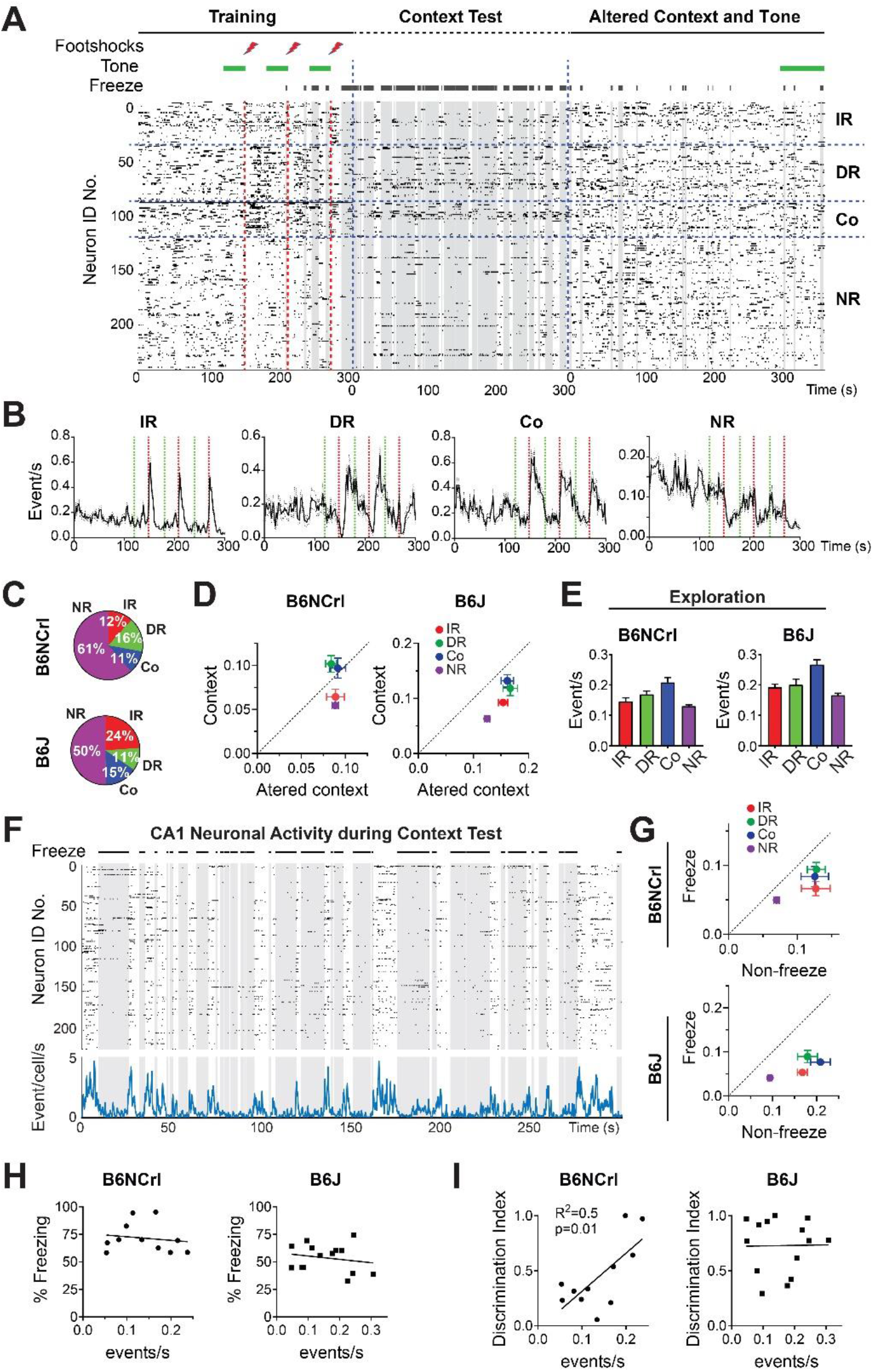
Neurons with delayed reactions play special roles in memory encoding. **(A, B)** Grouping neurons based on reactions to the US. **(A)** Raster plot of the CA1 neuronal activity of a representative mouse (B6J) during fear conditioning training, context test and altered context test. Neurons were grouped into four groups according to their response to the foot-shocks (IR: immediate response; DR: delayed response, Co: continuously responding; NR: Non-responding). **(B)** Average event rate for neurons in each group in B6J mice. **(C)** The percentage of neurons in each group. There were significantly more neurons in IR group in B6J than in B6NCrl (510/2139 vs 197/1668, p<0.001, Chi-square test). **(D)** The averaged activities of neurons in each group in context and altered context tests. In B6NCrl mice, the neurons with delayed reactions, including the DR and Co neurons, were preferentially activated in the original context although the overall neuronal activity was low in the context. The DR and Co neurons also showed higher activity levels than the other groups in the context but not in the altered context. **(E)** Averaged activities of neurons in each group during the exploration phase in fear conditioning training (the first 2 minutes). **(F)** Negative correlation between CA1 neuronal activity and freezing. The raster plot and average event rate of a representative mouse (B6J) during context test were shown. **(G)** Averaged neuronal activities of neurons in each group during freezing or non-freezing episodes during context test. The DR and Co neurons were more active than the other neuronal groups during freezing. Non-Freezing vs Freezing *** for all groups, Wilcoxon matched-pairs signed rank test **(H, I)** Activity of DR neurons during exploration predicts memory generalization level but not the freezing level in B6NCrl. (**H**) Correlation between freezing level in context and activity of DR neurons during exploration. **(I)** Correlation between discrimination index and activity of DR neurons during exploration.

Next we compared the activities of these 4 groups in the different stages of training and tests. In B6NCrl mice, DR and Co neurons were activated preferentially in the context test rather than in the altered context despite that the overall neuronal activities were lower in the context (Fig. 3D, left panel). Furthermore, DR and Co neurons showed higher activities than the IR and Co groups in the context test but not in the altered context. The preferential activation of DR and Co neurons in the original context but not in the altered context suggests these neurons play a special role in representing contextual fear memory. In B6J mice, the DR and Co neurons also showed higher activity level than IR and NR neurons in the original context, but the difference was less pronounced than in B6NCrl (Fig. 3D, left panel).

Consistent with a lack of locomotion during freezing, CA1 neuronal activities reduced significantly overall during freezing (Fig. 3F). We reasoned that if the memory represented by the hippocampus was driving the freezing behavior, then hippocampal neuronal activity during freezing would be associated with some on-going memory retrieval process. We analyzed the CA1 neuronal activity during non-freezing and freezing episodes during the context test. Despite the greatly reduced overall firing rate during freezing, DR and Co neurons maintained relatively high activities compared with IR and NR neurons in both sub-strains of mice (Fig. 3G).

Therefore, in B6NCrl mice, the neurons with delayed reaction showed 1) preferential activation in the original context; 2) higher activity levels than other neurons in the original context but not in the altered context, and 3) higher activity level during freezing. These all support a special role of these neurons in memory representation. An alternative explanation is that these neurons may be associated with fear or freezing itself. If so, these neurons would have lower activity than the other groups during the first 2 minutes of training, in which mice were actively exploring the context without fear-related freezing. But this was not the case (Fig. 3E). In B6J mice, DR and Co neurons showed a similar trend of preferential activation in the context and during freezing compared to other neurons, but the difference was less dramatic.

If neurons showing delay-phase responses participated in memory encoding, some of their activity features would be related to memory-guided behaviors. Indeed, in B6NCrl mice the activities of the DR neurons during the 2-minute exploration were positively correlated with the discrimination index but not the freezing level in Context (Fig. 3 H-I), indicating that the more active these neurons were during learning, the less generalized the memory was.

### Functional coupling of neurons in memory encoding and generalization

A proposed neuronal mechanism for learning and memory is experience-driven formation of neuronal assemblies. To examine functional coupling of neurons in the hippocampus we used the cell assembly detection (CAD) method (Russo and Durstewitz, 2017). CAD can detect assembly patterns across multiple timescales and time lags. In this method, cell pairs are tested for statistically significant correlations. We extended the correlation test to analyze the sparse neuronal activities recorded with calcium imaging. We then applied the extended method to extract neuronal pairs from whole imaging sessions including the fear conditioning training, context test and the altered context test (Fig. 4).

**Fig. 4:**
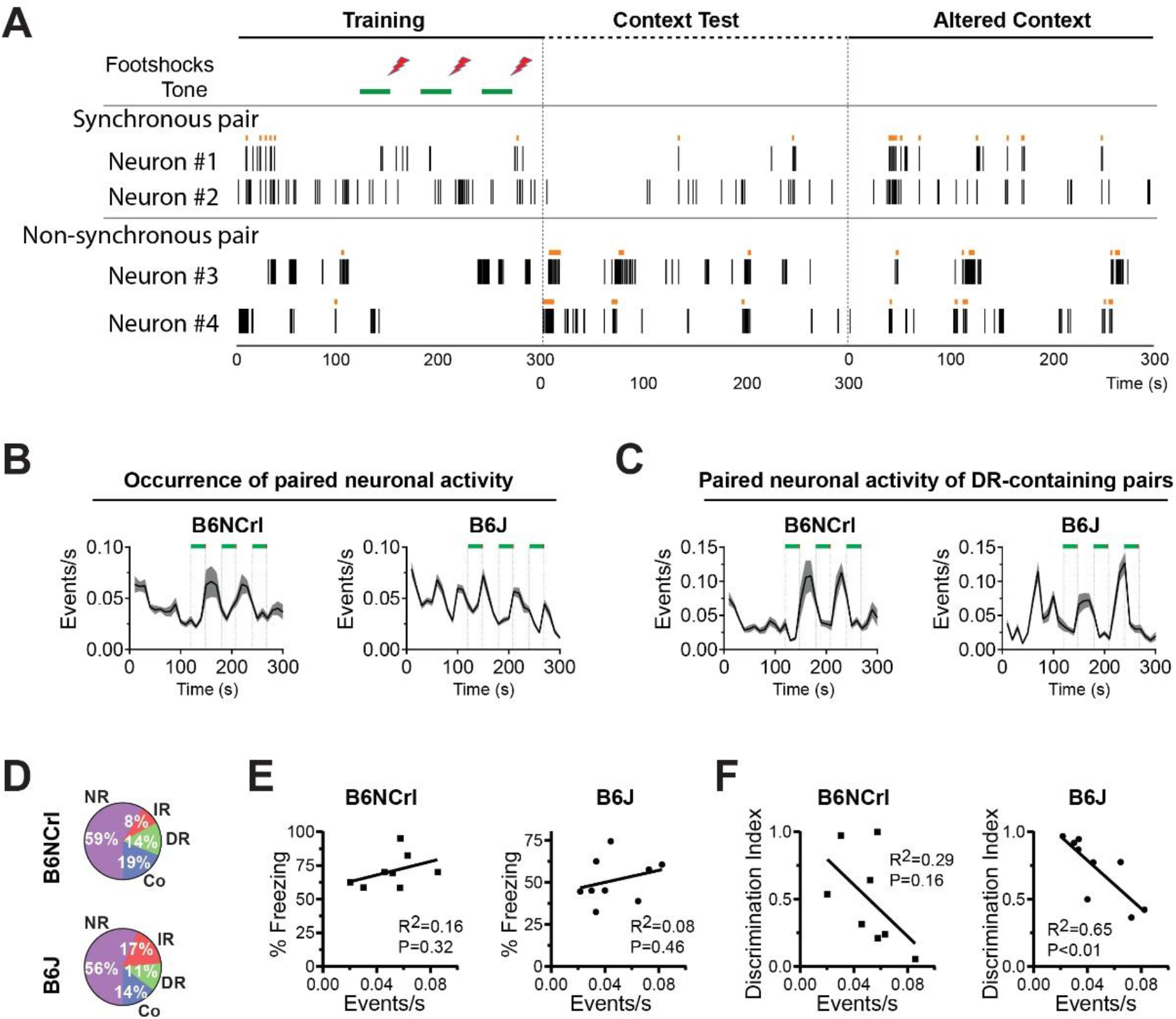
Coordinated neuronal activities of neurons with delayed reactions. **(A)** Examples of a synchronous neuronal pair (top) and a non-synchronous pair (bottom) from a representative mouse. The identified paired activities are marked with orange dots on top of the raster plots of calcium events. **(B)** The occurrence of all paired neuronal activities in fear conditioning training. The paired neuronal activities were averaged in 10-s bins. The data are mean ± SEM with the shaded area showing SEM (n= 8 B6Ncrl and 9 B6J mice). **(C)** The occurrence of paired neuronal activities of the neuronal pairs containing DR neurons in fear conditioning training. (**D**) The percentage of neurons paired with a DR neuron in the DR neuron-containing neuronal pairs. (**E**) The occurrence of DR neurons-containing neuronal pairs in training (the X axis) did not correlate with freezing in the Context in either sub-strain. (**F**) The occurrence of DR neurons-containing neuronal pairs in training (the X axis) correlated with discrimination index in B6J but not in B6NCrl.

We detected two types of neuronal pairs: synchronous neuronal pairs whose co-activity patterns happened in the same time bin, and nonsynchronous pair assemblies whose co-activity patterns happened with a time lag. Figure 4A shows examples for synchronous and nonsynchronous neuronal pairs detected from one mouse. We found that the detected pairs could occur inside each of the IR, DR, Co, or NR neuronal groups, or between two of these groups. We were able to detect the neuronal pairs in 9 B6J and 8 B6NCrl mice. Both sub-strains showed similar trends in the occurrence of paired neuronal activities (including all pairs identified): occurrence started high at the beginning of fear conditioning when the mice were first placed in the novel context, reduced when the mice became more familiar with the context, and increased again after the delivery of the US (Fig. 4B). These trends were largely parallel to the overall activities of the CA1 neurons in fear conditioning training (Fig. 2A). Due to the special roles of the DR neurons in memory representation, we examined the DR-containing neuronal pairs. While in B6NCrl the occurrence of paired neuronal activity of DR-containing neuronal pairs was similar to that of all neuronal pairs, in B6J this paired activity was low at the beginning of training and peaked at around 70 seconds (Fig. 4C), suggesting that the neurons in these DR pairs may become coupled in the process of learning. The neurons forming the DR-containing pairs came from all four groups, with NR neurons being the majority (Fi. 4D).

We then checked if the presence of neuronal pairs was related to the mice’s behavioral responses. In both sub-strains we did not find a parameter of the paired neuronal activities that correlated with freezing in the Context (Fig. 4E). However, we discovered that neuronal pairs containing DR neurons played a special role in generalization in B6J mice: the frequency of their paired neuronal activity during training was closely linked to the discrimination index (Fig. 4F). Specifically, the higher the paired neuronal activities were during training, the more generalized the contextual memories were, suggesting that functional coupling of the DR neurons with some other CA1 neurons is critical for memory generalization. Together with the above results (Fig. 3), these data demonstrate DR neurons play a unique role in memory representation in both sub-strains and that distinct neuronal activity features during learning in the two sub-strains predict the memory generalization levels tested later respectively.

### The major excitatory inputs to CA1 were similar in the two sub-strains

The above results indicate that neuronal reactions to the US are critically involved in memory encoding. What then are the cellular or circuitry mechanisms for the distinct reactions to US in the two sub-strains? We reason that the different reactions may come from 1) different excitatory inputs to CA1; 2) different excitability of CA1 neurons; and 3) the different inhibitory regulations. We tested these possibilities with calcium imaging and brain slice electrophysiology.

CA1 pyramidal cells receive their major excitatory inputs from the entorhinal cortex (EC) and the hippocampal CA3 region. To record the activity of synaptic inputs to CA1 in behaving mice, we injected AAVs mediating the expression of GCaMP6f fused with synaptic protein synaptobrevin-2 (Syb2-GCaMP6f) into either the EC or the CA3. Fusion to Syb2 enriched GCaMP6f to CA1 presynaptic terminals (Guo et al., manuscript under submission). We implanted GRIN lens into stratum lacunosum moleculare (s.l.m) or striatum radiatum (s.r.) layer in the CA1 to image the calcium signal in the synaptic terminals from EC or CA3, respectively (Fig 5A). The collective calcium signals from synapses were imaged and analyzed. The fluctuations of calcium signal in either EC or CA3 inputs correlated with the mouse’s locomotion (Fig. 5A, Supplementary Video-2). In both sub-strains the EC and CA3 input displayed strong responses temporally locked to each footshock (Fig. 5B-C). In addition, in both sub-strains the inputs from the EC or CA3 were significantly stronger during non-freezing episodes than during freezing episodes in context test (Fig. 5D). These results indicate that the synaptic inputs of EC or CA3 act similarly in B6J and B6NCrl, especially upon the delivery of the US, and the lack of the early-phase reaction in B6NCrl cannot be explained by the difference in these inputs.

**Fig. 5:**
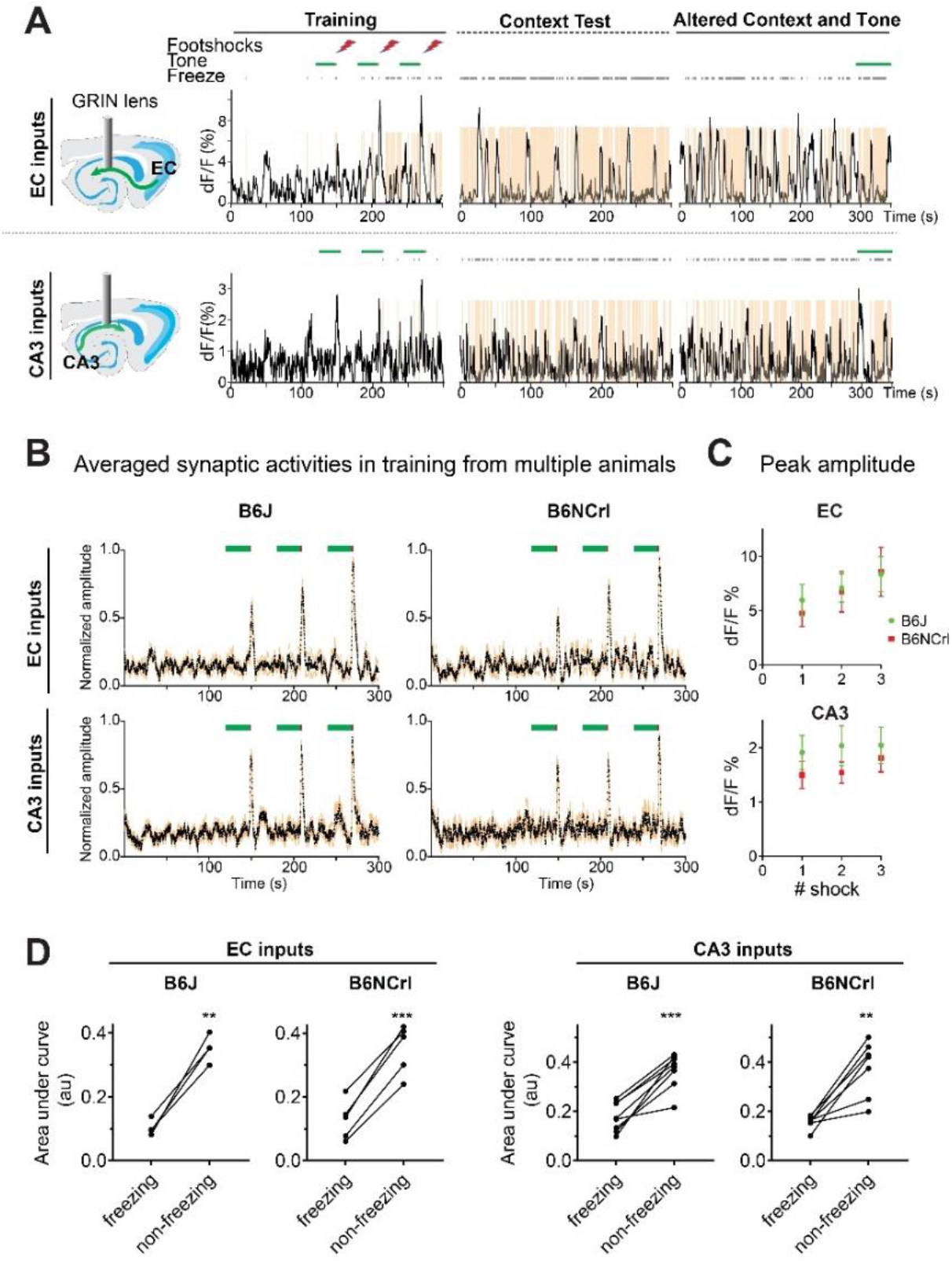
Activities of excitatory inputs to CA1 in learning and memory. **(A)** Schematics and example traces of calcium imaging of the collective activity of synaptic terminals in CA1 originating from the EC or the CA3, respectively, during fear conditioning training and tests. The black bars and the brown color shades indicate the time mice froze. **(B, C)** Synaptic terminals originating from EC and CA3 in both sub-strains showed peak activities upon the delivery of the US. (**B**) Averaged calcium signal of synaptic terminals in CA1 originating from the EC or the CA3, respectively. Data were normalized to maximum dF/F value in each animal before averaging the data from the recorded mice. The data are mean ± SEM (SEM shown by orange dots, n= 6 B6J mice and n= 6 B6NCrl mice for EC inputs. n= 8 B6J mice and n= 7 B6NCrl for CA3 inputs). **(C)** Quantification of peak activities upon each of the foot-shocks show no significant difference between B6J and B6NCrl. **(D)** Quantification of synaptic activities during freezing and non-freezing episodes. Synaptic activities were significantly stronger during non-freezing episodes than during freezing (** P<0.01, *** P<0.001 paired t-test).

### Inhibitory synaptic activities were distinct in B6J and B6NCrl

We further studied the intrinsic properties of the hippocampal circuits by comparing the excitability and synaptic activities of CA1 pyramidal cells in acute brain slices. The CA1 pyramidal cells in both sub-strains fired similarly in response to current injections, indicating similar levels of neuronal excitability (Fig. 6A).

**Fig. 6:**
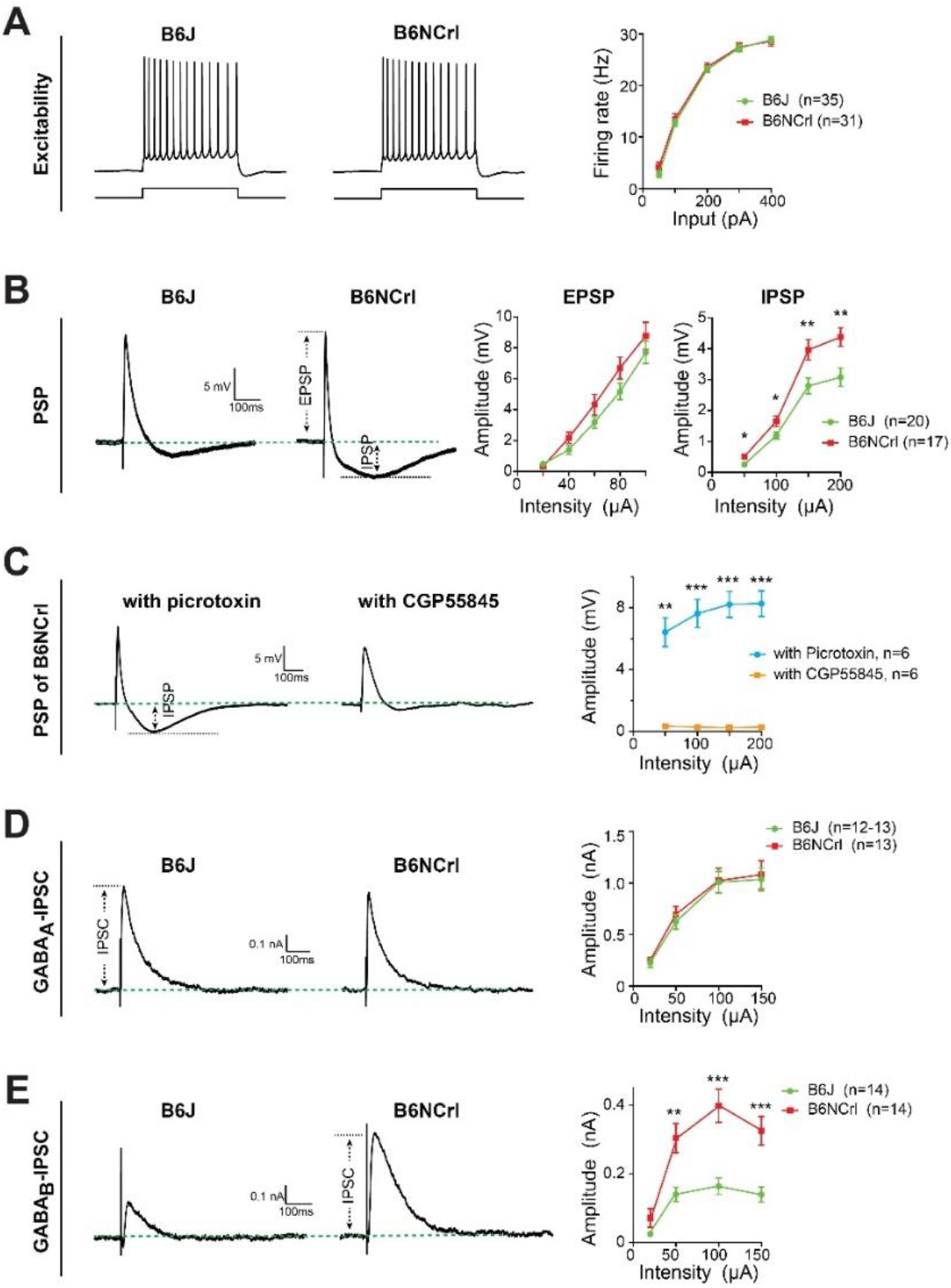
Distinct inhibitory synaptic activities in B6J and B6NCrl. **(A)** CA1 pyramidal cells of B6J and B6NCrl had similar excitability. Left: representative traces of action potentials induced by current injections in each sub-strain. Right: input-output relationship for induction of action potentials. **(B)** Electric stimuli-evoked post-synaptic potentials (PSPs) in B6J and B6NCrl. Left: representative traces of PSPs in CA1 pyramidal cells in response to electric stimulation at s.r.. Right: quantification of EPSP and IPSP components of the PSPs. The amplitudes of IPSPs from B6NCrl mice were significantly higher than those from B6J mice. **(C)** The IPSP component of the PSP was mainly mediated by GABA_B_ receptors. Left: representative traces of PSPs recorded in the presence of picrotoxin to block GABA_A_ receptors or CPG55845 to block GABA_B_ receptors, respectively. Right: quantification of IPSP amplitudes in the presence of picrotoxin or CPG55845. **(D)** Representative traces and quantification of GABA_A_ receptor-mediated IPSCs recorded in the presence of CPG55845. **(E)** Representative traces and quantification of GABA_B_ receptor-mediated IPSCs recorded in the presence of picrotoxin. The data in bar graphs are mean ± SEM. * P,0.05, ** P<0.01, ***P<0.001, two-way ANOVA followed by Student’s t-test.

We next compared the evoked synaptic responses in CA1. Electrical stimuli were delivered to s.r. and post-synaptic potentials (PSPs) were recorded in CA1 pyramidal cells in whole-cell configuration. Single electrical pulse induced bi-phasic PSPs--an EPSP phase followed by a prolonged IPSP phase (Fig. 6B). While the EPSP phase was similar between the two sub-strains, the amplitude of the IPSP phase was significantly higher in B6NCrl (Fig. 6B). The major mediator of IPSPs in CA1 are two types of GABA receptors--the ionotropic GABA_A_ receptors and the metabotropic GABA_B_ receptors. A selective GABA_B_ receptor antagonist, CGP55845,, massively reduced the IPSP phase, indicating that GABA_B_ receptors are the major mediator of this IPSP phase, and that GABA_B_ receptors-mediated synaptic inhibition is a major difference between B6J and B6NCrl (Fig. 6C). To further confirm, we selectively recorded GABA_A_ or GABA_B_ receptors-mediated IPSCs in the presence of selective antagonist for each of the receptors, respectively. There was no significant difference in GABA_A_ receptor-mediated IPSCs (Fig. 6D). In contrast, GABA_B_ receptors-mediated IPSCs in B6NCrl were significantly stronger than those in B6J (Fig. 6E). Together, the electrophysiological analysis shows that CA1 pyramidal cells in the two sub-strains are similar in intrinsic excitability; but they receive stronger GABA_B_ receptor-mediated synaptic inhibition in B6NCrl than in B6J.

## Discussion

In this study, we compared two closely related mouse sub-strains in learning and memory. Behaviorally, B6NCrl mice learn faster, freeze more and show more generalized fear memory. In neuronal activities, we saw distinct CA1 reactions in the two sub-strains following the US. Excitingly, we found neuronal signatures of generalization of contextual memories in both sub-strains, and intriguingly these signature activities between the sub-strains are not a difference in a continuous spectrum but rather distinct features. Furthermore, we observed different GABA_B_-receptor-mediated synaptic inhibition in the two sub-strains. How can we connect these dots to understand the neuronal mechanisms behind learning, memory and generalization?

### 1) Different feedforward inhibition reveals bi-phasic reactions to the US

What causes the early-phase reaction? Consistent with previous imaging and *in vivo* electrophysiological recordings (Jimenez et al., 2020; Lovett-Barron et al., 2014; Munera et al., 2001), one of the most obvious hippocampal activities during fear conditioning occurs upon the delivery of the US. Importantly, only by comparing the two sub-strains and observing that the early-phase is largely absent in B6NCrl, can we reveal that there are actually two phases of reactions. The hippocampus is a brain region where sensory information and their spatial/temporal relationships are integrated(Aronov et al., 2017; Manns et al., 2007). The hippocampus has access to multimodal sensory information (Eichenbaum et al., 1987; Fried et al., 1997; Sakurai, 2002). The US, footshocks themselves, and the vigorous escaping behaviors following footshocks may trigger a strong wave of sensory information to the hippocampus, which can trigger the early-phase reactions.

Why is the early-phase largely absent in B6NCrl? The hippocampus hosts diverse GABAergic inhibitory interneurons (Pelkey et al., 2017). The excitatory inputs from the entorhinal cortex, CA3 or thalamus activate these local interneurons to elicit feedforward inhibition of the principal cells— the pyramidal cells at CA1. The feedforward inhibition sets a brake on the activities of the postsynaptic neurons and restricts their firing to precise time windows (Pouille and Scanziani, 2001). A strong feedforward inhibition may minimize the activation of the pyramidal cells by the US (Fanselow et al., 1993; Lovett-Barron et al., 2014). Consistent with this idea, the GABA_B_ receptors-mediated inhibitory synaptic currents are significantly stronger in B6NCrl, while the intrinsic excitability and the excitatory synapses do not differ in the two sub-strains.

What causes the late-phase reaction? The late-phase is when the mice showed reduced locomotion (Fig. 2E). This “awake resting” period could be a time window for the neuronal representation of recent experience, especially emotionally charged experience, to be reactivated (Joo and Frank, 2018; Wu et al., 2017). This reactivation normally occurs in sharp-wave ripples, which are short periods of highly synchronous neuronal activity that originate from the CA3 and propagate to the CA1. Besides, the hippocampus is regulated by neuromodulatory systems, such as dopamine, acetylcholine, norepinephrine and serotonin (Avery and Krichmar, 2017). Some of them are released upon the delivery of the US (Jing et al., 2020; Robinson et al., 2019). The neuromodulatory systems normally produce effects at slower time scales (from seconds to minutes) (Parikh et al., 2007). It is possible that the late-phase arises from the slower kinetics of neuromodulation. Neuromodulators can increase CA1 neuronal activity by facilitating excitatory synapses or increasing neuronal excitability. They can also facilitate the “replay” of recent experience (McNamara et al., 2014). Selective pharmacological blockade of the specific neuromodulatory systems may shed more light on this in future.

What then are the functions of the two phases? Arising from different mechanisms, they may function differently. Through contextual fear conditioning, the emotionally neutral context becomes associated with the noxious US (Tronson et al., 2012). It’s generally believed that the formation of the memory of the context (the CS) occurs in the hippocampus while its association with US is in the amygdala (Fanselow and Poulos, 2005; Maren, 2001; Rudy et al., 2004). Therefore, the US does not necessarily have to be represented in the hippocampus; in fact, its presence in the hippocampus may be inhibited (Fanselow et al., 1993; Lovett-Barron et al., 2014). Consistent with this view, the mice learnt well both in the presence (B6J) and absence of the early-phase reactions (B6NCrl), suggesting that the representation of the US is not essential for contextual memory. However, a history of activation can influence the subsequent activation of hippocampal neurons for memory encoding (Cai et al., 2016). The activity in early-phase may contribute to the different behaviors, especially the different levels of generalization, in the two sub-strains, although the detailed mechanisms need further exploration. Unlike the early-phase, the late-phase may serve as a memory consolidation mechanism. Emotionally salient US may trigger replay of recent experience and a wave of release of neuromodulators, which either enhance ongoing neuronal representation of the context, or facilitate the replay (Singer and Frank, 2009). In either situation, this enhanced neuronal activity facilitates synaptic plasticity and memory strengthening. To further test this speculation, selective inhibition of the two phases with temporally precise optogenetics could be very informative.

### 2) Late-phase reaction in memory representation and generalization

Individual hippocampal neurons fire sparse spikes, but a high percentage of the cells fire within a few minutes of explorative behavior (Ahmed and Mehta, 2009). Do all the cells firing in a fear conditioning training equally participate in representing the context? Our results indicate that the neurons activating in the delayed phase, which account for 20-30% of the detected neurons, are preferentially engaged in contextual representation for 3 reasons. 1) They are preferentially “reactivated” when memory is retrieved. 2) They appear more active than the rest of the neurons when the mice freeze. 3) Activity of these neurons during training predicts the behavior responses, especially generalization, tested later. The predictive value of these neuronal activity features strongly supports their role in encoding the contextual memories.

Why are the neurons that are active in the late-phase particularly important? As discussed above, the activity in this phase may reflect a memory consolidation process. Contextual memory is first encoded in the exploration stage before the US. At this stage, this contextual memory may be weak and fragile. However, in the delayed phase, this weak contextual memory gets reactivated and strengthened by the neuromodulatory systems which are evoked by emotionally salient US.

The level of generalization may be determined at either memory encoding or retrieval stages (Asok et al., 2018; Berens and Bird, 2017; Kumaran and McClelland, 2012). Here we identified neuronal activity features in training that could predict generalization. In B6NCrl, the activity of DR neurons in exploration positively correlates with discrimination index. This is consistent with our previous work suggesting that increasing the number of CA1 neurons in memory encoding helps to generate a more precise and less generalized memory (also consistent with this notion, B6J has overall higher activity and less generalized memory compared to B6NCrl). In B6J, the activity of DR neuron-containing neuronal pairs negatively correlates with discrimination. This suggests that pairing of memory encoding neurons (the DR neurons) with some special CA1 neurons actively generalizes memory. We notice that the cells paired with DR neurons are mainly in the NR groups, indicating a coupling of memory-encoding and non-encoding neurons in the hippocampus. Identifying these cells may help us to elucidate the neuronal mechanisms of generalization. The two distinct activity features reflect different neuronal codes used by the mice for contextual memory and generalization.

Interestingly, in both sub-strains, we did not find an activity feature correlating with the freezing level in the original context. It is possible that certain high-dimension interactions among the neurons unidentified by our analysis are related to the freezing level. More likely, the freezing level is not determined by hippocampal memory system. Instead, other brain regions, such as the amygdala, hypothalamus or midbrain structures, may determine the freezing level based on the memory information from the hippocampus.

In summary, the CA1 reacts to the US in two phases. The early-phase may be triggered by the sensation related to the delivery of the US and is susceptible to GABA_B_-receptor-mediated feedforward inhibition; and the late-phase appears to be particularly important for memory encoding and generalization. The distinct neuronal activity features predicting generalization levels indicate that simple changes in hippocampal network properties can lead to diverse memory encoding mechanisms. The results also imply that hippocampal activity features in learning may be used as biomarkers in clinical settings to measure the risk of or vulnerability to mental disorders that involve overgeneralization of fear memories, such as post-traumatic stress disorder, panic disorders and anxiety disorders.

## Acknowledgments

This study was supported by Klingenstein-Simons Fellowship Awards in Neuroscience (to WX) and grant from NIH/NINDS (NS104828 to WX). We thank Dr. Ying Li for editing this manuscript, Ms. Wenqin Rita Du and Ms. Elizabeth Li for preparing AAVs.

## Author contributions

WX supervised the research; JG, DT, AB and WX designed the research and conducted the experiments; DL provided support on in viv imaging technology. All authors participated in analyzing the results and writing the paper.

## Methods

### Mice

C57BL/6J mice were purchased from UT Southwestern breeding core. C57BL/6NCrl mice were purchased from Charles River Laboratories. Mice were housed with 5 mice per cage before surgery and 1 mouse per cage after surgery on a 12 hr light /12 hr dark cycle with ad libitum access to food and water. 8-12 weeks old male mice were used for surgeries. 3-4 weeks old male mice were used for electrophysiology recording. All animal procedures were approved by UT Southwestern Institutional Animal Care and Use Committee and complied with the Guide for the Care and Use of Laboratory Animals by the National Research Council.

### Contextual fear conditioning

Mice were handled in 1-min epochs every day for 5 days before the first behavior session. For the 2-day protocol, 4 contexts were constructed as either training or testing contexts. Context 1 had metal wall and metal grid on the floor for delivering foot-shocks. It was scented with 10% ethanol. Context 2 was the same as Context 1 except that it was scented with 5% vanilla. Paper walls with stripes were added to Context 2 to construct Context 3. A plastic, smooth and opaque floor was added to Context 3 to make Context 4. During training, each mouse was allowed to explore Context 1 without any stimulation for 120 s. A tone of 82-83 dB was presented 120 s after the mouse entered the context and lasted 30 s. The last 2 s of the tone was paired with an electric foot-shock (2 s, 0.8 mA). The tone-and-shock pair was repeated two more times with 30 s interval between the end of shock and next tone. The mouse was removed from the chamber 30 s after the end of last shock. On day 2, 24 hr after the training session, mice were randomly divided into 4 groups. Contextual fear memory or its generalization was assessed by placing the mouse into one of the 4 contexts for 300 s without any tone or shock. The percentage of time the mouse spent freezing was measured. For the 3-day protocol for imaging experiments, mice were trained in Context 1 with the same tone-and-shock paradigm. On day 2, the mice were tested in Context 1 for 300s. On day 3, the mice were tested in Context 4 (the altered context) for 300 s without tone and 60 s with the tone. The percentage of time that each mouse spent freezing was determined using FreezeView software (Version 2.26, Actimetrics Inc, Wilmette, IL, USA). The position of animal in each video frame was determined manually with custom MATLAB scripts. The center of head was used to calculate the instant speed of animal.

### *In vivo* calcium imaging

To image calcium activity from neurons in CA1 pyramidal layer, mice were unilaterally injected with 500 nl of AAV_DJ_-hSyn-GCaMP6s-P2A-NLS-dTomato at AP: −1.95, ML: 1.25, DV: 1.35. A GRIN lens of 500 μm in diameter was implanted after viral injection at AP: −1.95, ML: 1.25, DV: 1.25. The lens was fixed on skull with Metabond and dental cement and was protected with a PCR tube. 3-4 weeks later the PCR tube was removed, and the base of camera was attached to the skull with dental cement. For CA1 presynaptic inputs imaging, mice were unilaterally injected with 500 nL of AAV_DJ_-hSyn-Syb2-GCaMP6f and highly diluted AAV_DJ_-hSyn-tdTomato (50:1) at AP: −4.8, ML:3.5, DV: 2.6 (EC) or bilaterally at AP: −2.35, ML: +/−2.6, DV:2 (CA3). A GRIN lens was implanted at AP: −1.95, ML:1.35, DV: 1.5 on the top of SLM layer for EC input imaging or at AP: −1.95, ML:1.25, DV: 1.3 on the top of SR layer for CA3 input imaging. The procedure of endoscope installation was the same as described above (Barbera et al., 2016)

### Brain slice electrophysiology

Transverse slices of the dorsal hippocampus (300 μm) were prepared with a vibratome (Leica VT1200) in ice cold cutting solution containing (in mM): 2.5 KCl, 1.2 NaH_2_PO4, 26 NaHCO3, 10 D-glucose, 213 sucrose, 5 MgCl_2_, 0.5 CaCl_2_. The slices were incubated for 30 min in artificial cerebrospinal fluid (ACSF) containing (in mM): 124 NaCl, 5 KCl, 1.2 NaH_2_PO_4_, 26 NaHCO_3_, 10 D-glucose, 1.3 MgCl_2_, 2.5 CaCl_2_ at 32 °C and for at least 1 h at room temperature. The cutting solution and ACSF were adjusted to pH 7.3-7.4 and 290 - 300 mOsm and constantly aerated with 95% O_2_/5% CO_2_. Whole-cell patch clamp recording was performed in a recording chamber perfused (~1 ml/min) with oxygenated ACSF at 26-28 °C. The recording pipettes (2.5-4 MΩ) were filled with internal solution containing (in mM): 125 K-gluconate, 20 KCl, 4 Mg-ATP, 0.3 Na-GTP, 10 Na_2_-phosphocreatine, 0.5 EGTA, 10 HEPES, adjusted to pH 7.3-7.4 and 310 mOsm. For electrically stimulating SR or SLM fibers, a focal stimulating pipette (~ 1 MΩ) was filled with ACSF and placed at 50 um below the surface of the slice and ~ 500 μm away from the recorded cell. When Picrotoxin or CGP55845 was used, they were added to the ACSF and perfused to recording chamber.

### Calcium imaging data analysis

For CA1 pyramidal cells imaging, the rigid motion was corrected by NoRMCorre and the calcium signal and deconvoluted events were extracted and analyzed with the extended CNMF for microendoscopic data (CNMF-E) approach (Pnevmatikakis and Giovannucci, 2017; Zhou et al., 2018). The detected components were manually examined and components without neuronal morphology were excluded.

To divide neurons into 4 groups based on their response to footshock, the activities of neurons during 0 – 10 s (IR group), 10 – 30 (DR group) time window from the onset of footshock were concatenated with those during −10 – 0 s (baseline window) before the onset of foot shock. A surrogate distribution was generated by randomly assigning all calcium events to non-overlapping frames in the concatenated time window for 10000 times. The difference between the number of events in either IR or DR time window and the number of events during baseline window was calculated for all 10000 shuffles. The difference value of actual event is compared to the 95^th^ percentile in the surrogate distribution. If the difference value is larger than the 95^th^ percentile in the surrogate distribution, this neuron was considered as significantly responding neuron for IR or DR. If a neuron is significant for both IR and DR, it was assigned to Co group but not IR or DR. Neurons without significant response were grouped into NR.

For imaging calcium activities in the presynaptic terminals, averaged fluorescence intensity was measured. A dynamic background was used. The minimum intensity within a 10-s window was used as the background for the image frame right after this 10-s period. For the frames in the first 10 s in each session, the minimum intensity in the first 10 s was used as their background.

### Cell assembly analysis

for detecting cell assemblies, we extend the cell assembly detection (CAD) method (Russo and Durstewitz, 2017), and includes a correction for non-stationarity. In brief, cell pairs are tested for statistically significant correlations. For each pair, a null hypothesis can be formulated in terms of a pairwise correlation quantity which can be shown to satisfy an F-distribution. However, we found that the original pairwise correlation test was not reliable for sparse spike train data, such as our calcium imaging data. Specifically, the F-distribution with the degree of freedom suggested by the original method was no longer an accurate proxy for the distribution of the pairwise correlation quantity. Therefore, we extended this pairwise correlation test by proposing an adjustment for the degree of freedom based on the sparsity of the data. The resulting test yielded more accurate testing performance on synthetic spike train data. For details, see Truong, Phan Minh Duc, “Cell assembly detection in low firing-rate spike train data” (2020). Mathematics Theses and Dissertations. (https://scholar.smu.edu/hum_sci_mathematics_etds/8).

### Quantification and Statistical Analysis

Data are presented as mean ± standard error of the mean (SEM). Sample number (n) indicates the number of cells or mice in each experiment and is specified in the results or the figures. The methods for statistical analysis are described in the results or the figures.

